# IGF-1 Peptide Mimetic-functionalized Hydrogels Enhance MSC Survival and Immunomodulatory Activity

**DOI:** 10.1101/2024.06.27.600680

**Authors:** Xiaohong Tan, Liufang Jing, Mohammadjafar Hashemi, Sydney M. Neal, Munish C. Gupta, Jacob M. Buchowski, Lori A. Setton, Nathaniel Huebsch

**Affiliations:** Department of Biomedical Engineering; Department of Orthopedic Surgery; Department of Mechanical Engineering and Materials Science; The Institute of Materials Science & Engineering, Washington University in St. Louis, USA

**Keywords:** Alginate hydrogels, dual peptide targeting, human mesenchymal stem cells, Insulin-like growth factor, immune modulation, Intervertebral disc cells

## Abstract

Human mesenchymal stem cells (MSCs) have demonstrated promise when delivered to damaged tissue or tissue defects for their cytokine secretion and inflammation modulation behaviors that can promote repair. Insulin-like growth factor 1 (IGF-1) has been shown to augment MSCs’ viability and survival and promote their secretion of cytokines that signal to endogenous cells, in the treatment of myocardial infarction, wound healing, and age-related diseases. Biomaterial cell carriers can be functionalized with growth factor-mimetic peptides (i.e. IGF-1 mimicking peptides) to enhance MSC function while promoting cell retention and minimizing off-target effects seen with direct administration of soluble growth factors. Here, we functionalized alginate hydrogels with three distinct IGF-1 peptide mimetics and the integrin-binding peptide, cyclic RGD. One IGF-1 peptide mimetic (IGM-3) in combination with integrin ligand was found to activate Akt and ERK1/2 signaling and support survival of serum-deprived MSCs. MSCs encapsulated in alginate hydrogels that presented both IGM-3 and cRGD showed a significant reduction in pro-inflammatory cytokine secretion when challenged with interleukin-1β. Finally, MSCs cultured within the cRGD/IGM-3 hydrogels were able to blunt pro-inflammatory gene expression of human primary cells from degenerated intervertebral discs. These studies indicate the potential to leverage cell adhesive and IGF-1 growth factor peptide mimetics together to control therapeutic secretory behavior of MSCs.

**Significance Statement:** Insulin-like growth factor 1 (IGF-1) plays a multifaceted role in stem cell biology and may promote proliferation, survival, migration, and immunomodulation for MSCs. In this study, we functionalized alginate hydrogels with integrin-binding and IGF-1 peptide mimetics to investigate their impact on MSC function. Encapsulating MSCs in the dual peptide (cRGD/IGM-3) hydrogels enhanced their ability to reduce inflammatory cytokine production and promote anti-inflammatory gene expression in cells from degenerative human intervertebral discs exposed to proteins secreted by the MSC. This approach suggests a new way to retain and augment MSC functionality using IGF-1 peptide mimetics, offering an alternative to co-delivery of cells and high dose soluble growth factors for tissue repair and immune-system modulation.

## Introduction

Transplanted mesenchymal stem cells (MSCs) are attractive for regenerative medicine for their ability to induce repair and modulate inflammation in host tissue [1]. Preclinical and *in vitro* studies have shown that soluble factors secreted by MSCs enhance host cell proliferation and migration, while combating apoptosis and inflammation [2, 3]. *In vivo*, transplanted MSC actively and continuously secrete cytokines, growth factors and extracellular vesicles [4–6] that circumvent a need to deliver very high doses of individual growth factors and/or vesicles. This ability of transplanted MSC to secrete multiple factors allows these cells to modulate multiple host cell behaviors, ultimately leading to therapeutic benefits. For example, transplanted MSC can inhibit T-cell proliferation by secretion of transforming growth factor beta (TGF-β) and prostaglandin E2 (PGE2), while also supporting tissue repair and regeneration by promoting anti-inflammatory M2 macrophage polarization [7, 8].

A challenge in MSC therapy is that the intrinsic function of these cells alone is often insufficient for targeted disease treatment [9]. Growth factors can significantly enhance the survival of transplanted MSCs and also bolster their therapeutic functions (e.g. cytokine production and extracellular matrix biosynthesis [10]). Prior work has shown that priming MSCs with growth factors prior to transplantation helps dampen host immune cell activation by triggering MSC secretion of trophic factors such as insulin-like growth factor (IGF-1), platelet-derived growth factor (PDGF), TGF-β, and interleukin-6 (IL-6) [11]. IGF-1, in particular, has been shown to be well-suited in this regard. For example, virally-induced overexpression of IGF-1 in MSCs enhanced their survival under stress conditions (e.g. transplantation into ischemic tissue) by triggering PI3K/Akt signaling [12]. Similarly, pretreating bone marrow-derived MSCs with soluble IGF-1 before transplantation into infarcted rodent heart muscle reduced host-production of pro-inflammatory cytokines TNF-α, IL-1β, and IL-6 gene expression [13]. Finally, IGF-1 overexpression in human umbilical cord-derived MSC significantly enhanced renoprotective effects in gentamicin-induced acute kidney injury by improving renal function, reducing histological injury and apoptosis, and upregulating genes involved in anti-oxidation, anti-inflammatory responses, and cell migration [14].

Despite the potential benefits of IGF-1 and other growth factors in enhancing the therapeutic potential of MSC, direct administration of soluble growth factors has led to undesired, dangerous off-target effects [15]. In one widely appreciated example, diffusion of soluble bone morphogenic protein 2 (BMP-2) away from the target site has been linked to ectopic bone formation, inflammation and pain [16, 17]. Likewise, cells transplanted without a biomaterial carrier may also escape the target site and cause intended side effects and/or injury. For example, MSC transplanted into degenerative intervertebral discs (IVD) without a carrier escaped the transplant site and subsequently triggered osteophyte formation [16, 18].To mitigate these issues, biomaterial carriers have been developed to entrap cells, and bioactive peptide mimetics of growth factors have in turn been grafted to provide an immobilized source of growth factor signaling to these carriers [4, 19]. For example, peptides derived from BMP-2 have been grafted to alginate gels, and this has conferred an ability of these gels to induce osteogenesis in encapsulated MSC [19, 20]. Similarly, the C domain peptide of IGF-1 has been attached to chitosan gels, which enhanced transplanted MSC viability and improved cells’ ability to enhance host tissue repair [21]. Self-assembling peptide gels that incorporate the C-domain of IGF-1 have also been used to enhance MSC survival and augment the ability of transplanted MSC to induce anti-inflammatory macrophage polarization in the context of treating acute kidney injury [22].

Here, we sought to use IGF-1 peptide mimetics to enhance survival and immunomodulatory capabilities of MSC. In a screen of three candidate peptides, a bioactive IGF-1 peptide mimetic that was identified previously through phage display [23] proved much more potent than the C-domain of IGF-1 in improving MSC survival under serum-deprived culture conditions. Survival was improved markedly beyond what we observed in MSC cultured in gels presented with cell adhesive cyclic RGD (cRGD) peptides alone, suggesting a potential unique role for growth factor mimetics in promoting cell survival. Interestingly, although this new peptide does not share sequence similarity with IGF-1, it activated Akt signaling similar to the full-length growth factor, and significantly blunted the inflammatory response of MSC provoked with IL-1β. In co-culture studies, MSCs encapsulated within the alginate hydrogel functionalized with both the IGF-1 peptide mimetic and cRGD markedly blunted inflammatory responses in primary cells retrieved from degenerative human IVDs. Our findings indicate that the combination of cell-adhesive and IGF-1-mimicking peptides not only enhances MSC survival and the secretion of pro-reparative factors but also enhances the ability of these cells to attenuate inflammatory responses.

## Materials and Methods

### Peptide-modified Hydrogel

High-molecular-weight, high G-block-containing alginate (Manugel, Dupont, Wilmington, DE USA) was dissolved at 1% (w/v) solution in Dulbecco’s PBS (dPBS). Alginate polymer solution was purified by dialyzing against deionized water. For peptide conjugation to alginate, Alginate was modified with N-terminal cysteine presenting IGM peptides using maleimide-thiol click chemistry described in our prior study [24]. Briefly, to graft each of the cysteine-terminated IGM peptides (Fig. 1A, GenScript, Piscataway, NJ), the maleimide-modified alginate was dissolved at 1% w/v in ultrapure water for 2 h. Either IGM-1 (CGGGYGSSSRRAPQT, [21]), IGM-2 (CGGGSLDESFYDWFERQLGKK, [23]), or IGM-3 (CGGGDYKDLCQSWGVRIGWLAGLCPKK, [23]) was dissolved in ultrapure water and added dropwise to the maleimide-modified alginate solution at an equivalent to twice the molar ratio of BMPH conjugated to the alginate polymers [e.g., when the measured molar output for maleimide was 5 maleimide/polymer, a molar input of 10 IGM-polymer was used]. Alginate was modified with cRGD peptides coupled through a heterobifunctional spacer using Strain Promoted Azide Alkyne Cycloaddition (SPAAC) click chemistry, as described in our prior study [25]. Briefly, BCN-alginates were dissolved at 1% w/v in Dulbecco’s phosphate-buffered saline (dPBS) for 2 h. Azide cRGD (cyclo[Arg-Gly-Asp-D-Phe-Lys-(azide)]; Peptides International; Louisville, KE, USA) was then added dropwise to BCN-alginate solution at an equivalent to twice the molar ratio of BCN amine conjugated to the alginate polymer and allowed to react at room temperature with constant stirring for 24 h. Following dialysis (3400 Da cutoff) for 72 h, polymers were sterile filtered (0.42 µm) and freeze-dried polymers were maintained under aseptic conditions and stored at -80 °C for further use. Polymers prepared for subsequent studies were unmodified alginate, 10 µM cRGD-alginate, 50 µM cRGD-alginate, 50 µM cRGD/ 50 µM IGM-1, 50 µM cRGD/ 50 µM IGM-2, 50 µM cRGD/ 50 µM IGM-3, or 10 µM cRGD/ 50 µM IGM-3. Polymer solution was ionically crosslinked using divalent calcium ions and used for subsequent studies.

**Figure 1.**
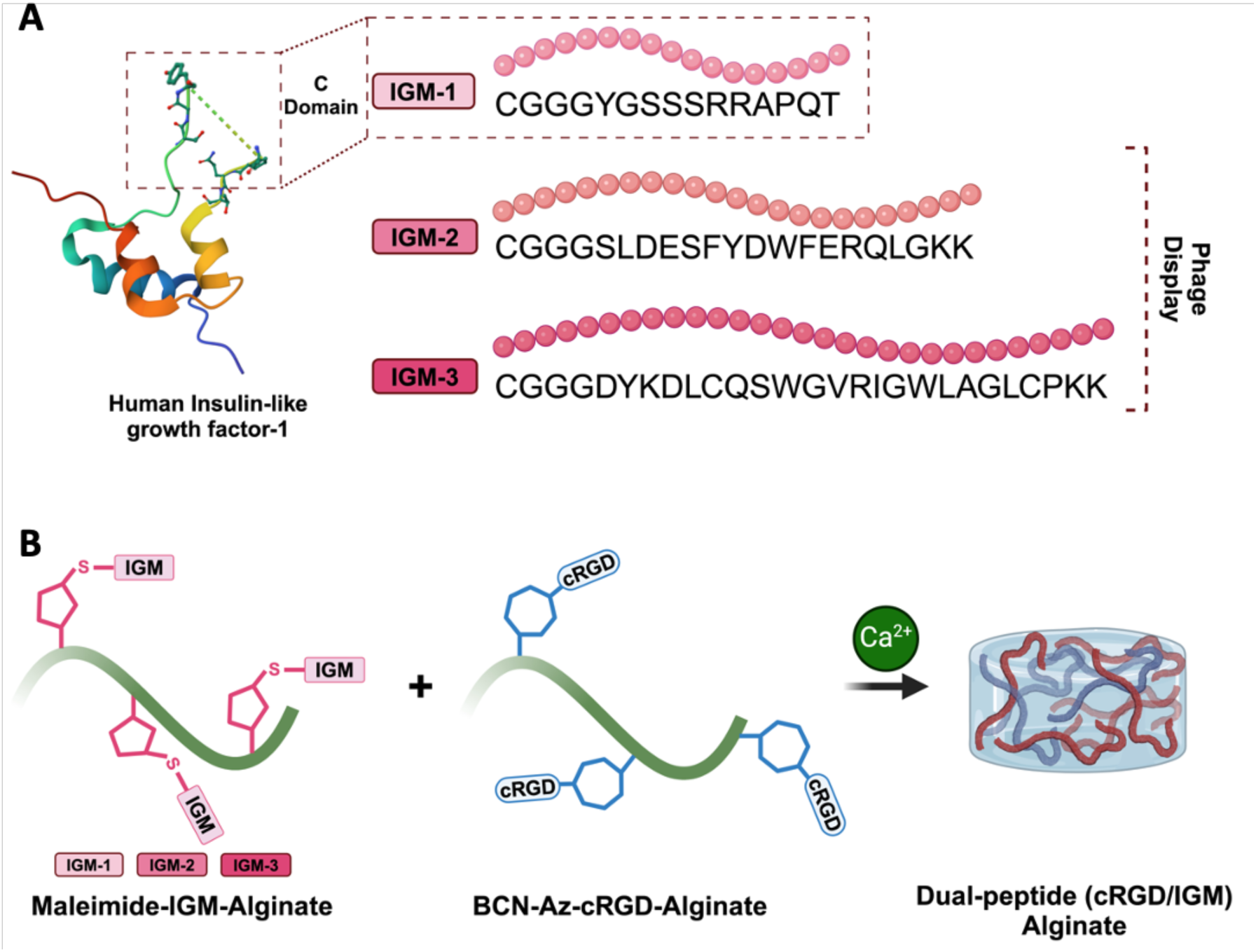
Schematic Representation of Bioconjugation Reactions Involving Insulin-like Growth Factor-1 Peptide Mimetics. A) Structural diagrams of Insulin-like growth factor peptide mimetic (IGM-1, IGM-2, IGM-3) as candidate molecules. IGM-1 represents the c-domain of the full-length IGF-1 [53]. In contrast, IGM-2 and IGM-3 are peptide mimetics that were identified through phage-display studies. B) Depiction of the conjugation techniques and hydrogel formation used in the study. Maleimide-thiol Michael-type addition reaction for IGM peptide conjugation and strain-promoted alkyne-azide cycloaddition for cRGD peptide conjugation are illustrated. Maleimide−IGM−alginate was combined with BCN−Az-cRGD−alginate at various peptide concentrations, and then ionically crosslinked with divalent calcium ions to form a combination peptide (cRGD/IGM) alginate hydrogel.

### Human Bone-marrow Derived Mesenchymal Stem Cell (MSC) Culture

Human bone marrow-derived MSC (RoosterBio^TM^, Frederick, MD) were acquired and expanded according to RoosterBio^TM^ expansion protocol. Briefly, MSCs were plated in tissue culture flasks in Roosterbasal^TM^ medium supplemented with RoosterBooster^TM^ until 80% confluence at 37 °C and 5% CO2. MSCs were detached using Gibco TrypLE^TM^ Express Enzyme (1X) (ThermoFisher Scientific, Waltham, MA) and passaged. Cells from passages 2 to 5 were used for experiments to ensure their multipotent characteristics.

### Cell Survival Assessment

Sterile, freeze-dried polymers were dissolved in 100 mM HEPES buffer (pH 7.2) at 2% w/v concentration and mixed overnight at 4 °C on a rotator. MSCs were prepared at a density of 2.5 x 10^6 cells/mL and mixed with an equal volume of the alginate solution to achieve a final concentration of 1% w/v alginate in the hydrogels. These hydrogels were then crosslinked using a calcium sulfate slurry to a final concentration of 100 mM calcium. The mixtures were quickly poured between two glass plates spaced 1 mm apart and left at room temperature for one hour to complete the crosslinking process. The formed hydrogels were punched into 5 mm discs using a biopsy puncher and placed in a 96-well plate containing RoosterBasal™ medium supplemented with RoosterBooster™ for 24 hours, then switched to RoosterBasal™ medium alone for 7 days at 37 °C and 5% CO2, with a 50% media change every 48 hours. Human recombinant insulin-like growth factor-1 (rhIGF-1, 500 ng/mL, Peprotech, Cranbury, NJ) was added to the unmodified alginate and cRGD-alginate hydrogel as positive controls.

Cell viability was assessed using the Resazurin-based Alamar Blue assay (Thermo Fisher Scientific). After culturing in RoosterBasal™ for 4 days, all hydrogel conditions encapsulated with MSCs were gently rinsed with phosphate-buffered saline (PBS). Fresh medium containing 10% Alamar Blue solution was then added to each well, and the plates were incubated for an additional 4 hours at 37°C in a humidified atmosphere with 5% CO₂. The fluorescence intensity of the medium was measured spectrofluorometrically at 560 nm/590 nm excitation/emission using an EnSight Multimode Plate Reader (PerkinElmer, Waltham, MA). The Resazurin readouts on day 7 were normalized to the readings obtained at 24 hours from the same gel, with each gel treated as a biological replicate.

### Signaling Pathway Activation Studies

Both non-phosphorylated and phosphorylated forms of the proteins AKT1/2/3, p-AKT 1/2/3 (pS473), total ERK 1/2, and p-ERK 1/2 (Thr202/Tyr204) were quantified using AlphaLISA SureFire Ultra Assay Kits (high volume, Revvity, Waltham, MA). For this purpose, sterile polymers, specifically 10 µM cRGD-alginate, and 10 µM cRGD/50 µM IGM-3-alginate, were dissolved at 1% w/v in sterile 100 mM HEPES buffer (pH 7.2) and stirred continuously overnight at 4 °C. Subsequently the next day, these solutions were rapidly mixed with sterile calcium sulfate slurry to achieve a final concentration of 100 mM calcium in the gel. The resulting mixtures were then poured between two glass plates separated by 1 mm spacers and allowed to crosslink at room temperature for one hour. Following crosslinking, the hydrogels were punched into 5 mm discs using a biopsy punch and transferred to wells of a 96-well plate containing RoosterBasal™ medium. These gels were incubated at 37 °C and 5% CO2, with the incubation medium refreshed six times at regular intervals over three days to remove excess calcium before cell seeding. Prior to the signaling assay and small molecule inhibition, the medium in the culture flasks containing MSCs was replaced with RoosterBasal™ without RoosterBooster™ supplementation for 24 hours. MSCs were then detached using Gibco TrypLE™ Express Enzyme (1X) (ThermoFisher Scientific), resuspended in RoosterBasal™, and seeded onto the gel at a density of 50,000 cells per well. After cell seeding, treatments were administered to designated wells, including AZ7550 Mesylate (20 µM, MedChem Express), GSK1904529A (3 µM, MedChem Express), LY294002 (10 µM, MedChem Express), U0126 (10 µM, MedChem Express), rhIGF-1 (500 ng/mL, Peprotech), or no treatment. For antibody blocking following signaling protein assessment, cells were resuspended in 2x antibody working solution of either mouse IgG1 isotype control (MAB002R, R&D Systems) or human/mouse IGF-I R /IGF1R antibody (MAB391, R&D Systems) prepared in basal medium supplemented with 0.1% BSA (1mg/mL), and pre-incubated on a rocker for 20 minutes at room temperature. After rocking, either recombinant human IGF-1 or basal medium alone was added to achieve a final antibody concentration of 500 ng/mL. For both small molecule inhibition and antibody blocking studies, cells were allowed to attach for 2 hours at 37 °C and 5% CO2, after which each cell-seeded hydrogel was considered an individual sample for subsequent analysis. The AlphaLISA assay was then performed according to the manufacturer’s protocol. Briefly, the medium was removed, and lysis buffer was added to each well. AlphaLISA reaction buffers (donor and acceptor mixes) were subsequently added to the lysate before the wells were read using an EnSpire Multimode Plate Reader (PerkinElmer) with the AlphaScreen protocol at an excitation of 680 nm and emission of 615 nm. Maximal protein phosphorylation was quantified by the ratio of phosphorylated protein to total protein in MSCs for p-AKT1/2/3 (pSer473) and p-ERK1/2 (pThr202/pTyr204). pAKT/total AKT of cells cultured on cRGD/IGM-3 was normalized to cRGD with the addition rhIGF-1 in studies quantifying the activation signal and antibody blocking study. In studies of small molecule inhibition, data was normalized to cRGD/IGM-3 to quantify the pAKT after inhibition.

### Multiplexed Secreted Protein Analyses

MSCs were encapsulated in each alginate gel as described in the Cell Survival Assessment Section. Following initial encapsulation, the hydrogel-containing cells were maintained in RoosterBasal™ medium and treated with 1 ng/mL of IL-1β (Sigma) for 4 days at 37 °C and 5% CO_2_. After the treatment period, cell culture supernatants were collected, transferred to tubes, and stored at -80°C until analysis. Protein levels of pro-inflammatory cytokines and chemokines in the supernatants were quantified using Luminex xMAP technology. The multiplex analysis was performed using a Luminex™ 200 system (Luminex, Austin, TX) by Eve Technologies Corp. (Calgary, Alberta, Canada) with human multiplex kits from MilliporeSigma (Burlington, Massachusetts, USA).

### Primary Human Intervertebral Disc (IVD) Cell Culture

Intervertebral disc cells (n≥3, male and female, ages 40 to 70) were isolated from the nucleus pulposus region of to-be-discarded surgical waste tissues; patients were de-identified and only sex, race and age of the patient were recorded (non-human subjects research exempt from pathology review; approved by Washington University IRB). For these studies, disc tissue was isolated from 6 patients (3 male, 3 female) aged 40-70. Disc cells were isolated as described in our previous studies [24]. Briefly, tissue fractions were removed from surgical samples and digested for 4 h at 37°C and 5% CO_2_ in medium containing 0.4% type II collagenase. (Worthington Biochemical, Lakewood, NJ and 2% pronase (Roche, Basel, Switzerland). Cells were passed through a 70 μm cell strainer (Thermo Fisher Scientific, Waltham, MA). The isolated cells (P0) were plated in tissue culture flasks in Ham’s F12 medium (Thermo Fisher Scientific) supplemented with 10% fetal bovine serum (FBS) and 1% penicillin-streptomycin (P/S) until confluence. For co-culture experiments, primary human disc cells were seeded in the bottom wells of 24-well Corning^TM^ Transwell^TM^ plates (Corning, NY, USA) with 0.4 μm permeable polyester membrane inserts. These cells were cultured at a density of 30,000 cells per well in F12 medium containing 10% FBS and 1% penicillin-streptomycin under hypoxic condition (37°C and 5% CO_2_ and 2% O_2_) for 24 h.

### *In Vitro* Transwell Co-culture of Monolayer IVD cells and MSCs Encapsulated in Hydrogel

At 24 hours, the encapsulation of bone marrow-derived MSCs was performed as previously described. MSCs were encapsulated within alginate, 10 µM-cRGD, or 10 µM-cRGD/50 µM-IGM-3 alginate hydrogels. The Transwell inserts were pre-moistened with RoosterBasal™ medium, and each hydrogel was then placed within a Transwell insert. These inserts were subsequently positioned in wells containing primary human disc cells, which were cultured in RoosterBasal™ medium treated with 1 ng/mL IL-1β under hypoxic (2% O_2_) conditions to challenge both the MSC and IVD cells. Monolayer IVD cells with no co-culture treated with 1 ng/mL IL-1β served as control. All wells were incubated for 4 days, with 50% media changes conducted at 48-hour intervals.

### Cell Lysate Collection and RNA Processing

Following the 4-day incubation period, hydrogels containing MSCs were removed from the Transwell inserts. IVD cells that were attached to the well were washed with 1% dPBS and lysed using RLT buffer (Qiagen, Hilden, Germany) and 1% mercaptoethanol (Sigma). mRNA isolation was performed using the RNeasy Kit with DNase I digestion (Qiagen Iberia, Madrid, Spain), following the manufacturer’s protocol. Briefly, NP cell samples were homogenized using a QIAshredder™ column, processed through an RNeasy spin column for RNA binding, followed by washing, and DNA digestion with DNase I. The purified RNA was eluted after two washes with washing buffer. The quality and quantity of the isolated mRNA were assessed by measuring the absorbance ratio at 260/280 nm using a NanoDrop One Spectrophotometer (Thermo Fisher Scientific).

### cDNA Library Generation and Bulk RNA-sequencing (bulk RNA-seq) with Data Standardization

The input samples were submitted to the Washington University Genome Technology Access Center to obtain and sequence the cDNA libraries (Illumina NovaSeq X Plus) according to manufacturer’s protocol. Briefly, RNA-seq reads were aligned to the Ensembl release 101 primary assembly with STAR version 2.7.9a. Gene counts were derived from the number of uniquely aligned unambiguous reads by Subread: feature Count version 2.0.3. Isoform expression of known Ensembl transcripts were quantified with Salmon version 1.5.2. Sequencing performance was assessed for the total number of aligned reads, total number of uniquely aligned reads, and features detected. The ribosomal fraction, known junction saturation, and read distribution over known gene models were quantified with RSeQC version 4.0.

All gene counts were imported into the R/Bioconductor package EdgeR and TMM normalization size factors were calculated to adjust samples for differences in library size. Ribosomal genes and genes not expressed in the smallest group size minus one samples greater than one count-per-million were excluded from further analysis. The TMM size factors and the matrix of counts were then imported into the R/Bioconductor package Limma. Counts-per-million (CPM) normalization was applied to the raw gene expression counts to account for differences in sequencing depth between samples. For each gene, the raw counts were divided by the total number of reads in the sample and then multiplied by one million. To correct for potential batch effects, Surrogate Variable Analysis (SVA) was performed using the sva package in R [26]. Surrogate variables were identified using the sva function which estimated the number of hidden sources of variation. Once the number of surrogate variables was determined, sva function was applied to the normalized count matrix to account for batch effects. The final linear model included both the treatment conditions (monolayer and hydrogel groups) and the surrogate variables as covariates. Weighted likelihoods based on the observed mean-variance relationship of every gene and sample were then calculated for all samples, and the normalized count matrix was transformed to moderated log2 CPM with Limma’s voom With Quality Weights. Each cell in the matrix contains the log2-transformed CPM value for a given gene in a given sample. The performance of all genes was assessed with plots of the residual standard deviation of every gene to their average log-count with a robustly fitted trend line of the residuals. The data were first centered and scaled before PCA to ensure that each gene contributed equally to the analysis regardless of its expression range. PCA was conducted using the prcomp function in R (RStudio Team 2020) to reduce dimensionality of the data. The input data for PCA consisted of the log2 CPM matrix (rows correspond to genes and columns correspond to samples) generated from the voom With Quality Weights transformation in the Limma package. The first two principal components were plotted to assess the distribution of samples and potential clustering between different groups. Differential expression analysis was then performed to analyze for differences between conditions in NP cells, respectively. The results were filtered for only those genes with Benjamini-Hochberg false-discovery rate adjusted p-values less than or equal to 0.05 and an absolute log2 fold change equal to 2 or greater. Gene Ontology analysis was conducted for differentially expressed genes using the clusterProfiler package (v3.14.3) in R. Results were filtered for terms with an adjusted p-value less than or equal to 0.05 and log2 fold-change less than -2 or greater than 2.

### Statistical Analysis

Statistical analyses were performed using GraphPad Prism 9 (Graph Pad Software, Inc., San Diego, CA, USA). Significance was determined by one-way ANOVA with Tukey’s post hoc test (P < 0.05). Two-tailed Student’s t-tests were used to estimate statistically significant differences in Akt phosphorylation between 10 μM cRGD-alginate supplemented with rhIGF-1 and 10 μM cRGD / 50 μM IGM-3-alginate combination. Data represent the mean ± SD. Different letter denotes statistical significance.

## Results

### Effects of Growth Factor-Mimicking Peptide Grafted Alginate on Encapsulated MSCs

We first screened three candidate peptides that either mimic IGF-1 or the structurally similar growth factor, insulin (Fig. 1A). These peptides were previously identified based on their sequence similarity to IGF-1 or insulin (IGM-1, [21]) or through phage-display studies (IGM-2, IGM-3, [23]). For screening studies, alginate polymers were grafted with IGF-1 peptide mimetics or cRGD using coupling chemistry described previously (Fig. 1B, [24, 25]). MSCs were encapsulated into alginate gels via calcium crosslinking. Among the three peptides, IGM-3 best maintained MSC viability in serum-deprived media, with cell viability approaching levels that were observed in cRGD-coupled alginate gels supplemented with a high dose of soluble recombinant human IGF-1 (rhIGF-1 at a dose of 500ng/mL, Fig. 2A). In contrast, alginate gels coupled with cRGD together with either IGM-1 or IGM-2 elicited only a modest improvement in MSC viability over what was achieved with cRGD-coupled alginate alone. Optimizing co-presentation of cRGD together with IGM-3 further enhanced material performance, to the point that cRGD/IGM-3 functionalized alginates elicited a significant improvement in MSC viability over what was achieved with cRGD and a high dose of soluble IGF-1 (Fig. 2B). Similarly, cRGD/IGM-3-alginate significantly supported MSC viability compared to IGM-3 hydrogel alone, which suggests the presentation of both cell-adhesive and IGM-3 peptide mimetic is necessary to support cell survival under no serum condition (Supp. Fig. 1).

**Figure 2:**
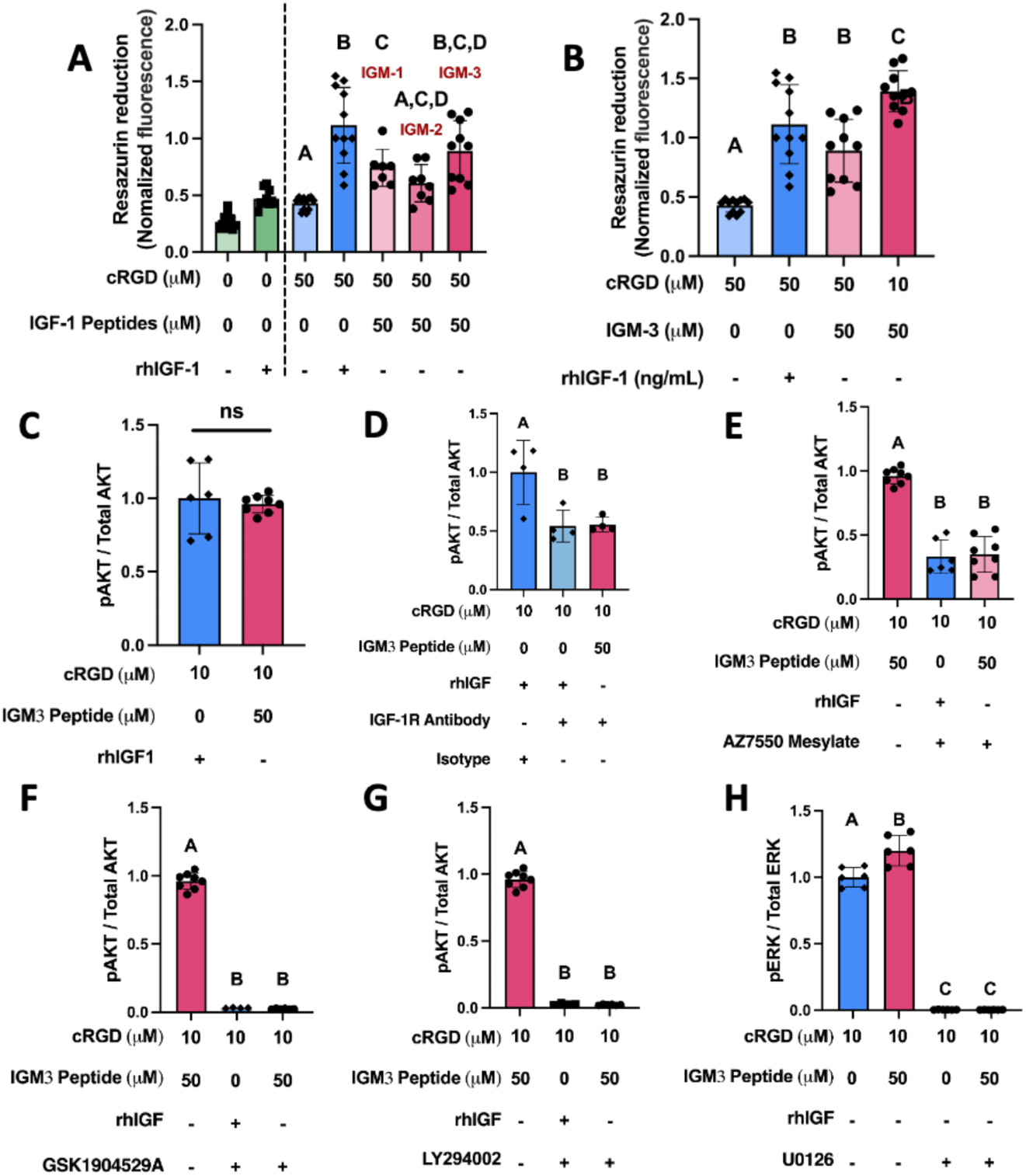
Mechanism of Action of cRGD/IGM-3 Alginate Hydrogel on MSCs. A) Resazurin reduction quantification in MSCs cultured in serum-free medium across different hydrogel formulations (alginate, alginate + rhIGF-1 [500 ng/mL], cRGD-alginate, cRGD-alginate + rhIGF-1, and alginate modified with IGM peptides [IGM-1, IGM-2, IGM-3] each combined with cRGD) on Day 7, normalized to Day 1 values. B) Comparison of resazurin reduction in MSCs encapsulated in cRGD-alginate, or varied peptide density of cRGD and IGM-3 on Day 7, normalized to Day 1 values. C) Akt phosphorylation levels (pAkt / Total Akt) measured two hours post-cell attachment on cRGD-alginate supplemented with rhIGF-1 (500 ng/mL), indicating activation similar to cRGD/IGM-3 alginate. D) IGF-1 receptor blocking of cRGD-alginate +rhIGF-1 or cRGD/IGM-3 demonstrates similar phosphorylation level. IGF-1 receptor small molecule inhibitor (AZ7550 Mesylate, 20μM) attenuated Akt phosphorylation to a greater extend compared to antibody blocking of IGF-1 receptor. E-F) Effect of IGF-1 receptor inhibitor (AZ7550 Mesylate, 20μM) and IGF-1 & insulin receptor inhibitor (GSK1904529A, 3μM) on Akt activity in hydrogels containing rhIGF-1 or IGM-3, demonstrating receptor-specific binding of IGM-3. G) Impact of PI3K/Akt pathway inhibition (LY294002, 10μM) on Akt phosphorylation in hydrogels supplemented with rhIGF-1 and cRGD/IGM-3 after 2 hours. H) ERK activity modulation by the ERK 1/2 pathway inhibitor (U0126, 10μM) in MSCs cultured on cRGD/IGM-3 alginate, showing pathway involvement in IGM3-mediated effects. Data points represent biological replicates with error bars indicating ±SD. Data normalized to cRGD-alginate +rhIGF-1 in C-H. Statistical analyses were performed using one-way ANOVA with Tukey’s multiple comparisons test; non-significant results are indicated by the same letter, while significant differences are marked with different letters at *p < 0.05*.

IGF-1 triggers pro-reparative behaviors (e.g. proliferation) and blocks apoptosis in MSC and other cell types (e.g. cardiomyocytes and skeletal myocytes) primarily by activating the PI3K/AKT [27, 28]. Since IGM-3 does not share a sequence with IGF-1, we questioned whether it could activate signaling in a similar manner. Thus, we probed the activation of the Akt pathway in MSCs by IGM-3. Surprisingly, short term culture in cRGD and IGM-3 peptide-coupled alginate activated Akt signaling in MSCs to levels similar to culture in cRGD-coupled alginate with the addition of a high dose of soluble IGF-1 (500ng/mL; Fig. 2C).

To determine if IGM-3 induced Akt signaling activation through the same receptor as soluble IGF-1, cells were pre-incubated with IGF-1 receptor antibody [29] and cultured atop cRGD-alginate supplemented with soluble IGF-1 or cRGD/IGM-3-alginate gels. Functional blocking of the receptor revealed that Akt activity significantly reduced suggesting IGM-3’s affinity to IGF-1 receptor (Fig. 2D). To further probe the mechanism of IGM-3 activation, cells were cultured on the cRGD/IGM-3 hydrogel in the presence of either AZ7550 Mesylate (an active metabolite of AZD9291, IGF-1 receptor inhibitor; [30]) or GSK1904529A (IGF-1 receptor and insulin receptor inhibitor; [31]). Akt activity was reduced in MSCs on cRGD/IGM-3-alginate treated with IGF-1 receptor inhibitor (AZ7550 mesylate), and decreased further when both receptors were inhibited to a level similar to that observed when incubated with LY294002, an inhibitor of all Akt activation ([32]; Fig. 2E-G). Dose response was performed with Akt inhibitor to ensure dosage used did not affect cell viability. Importantly, gel-grafted IGM-3 and soluble IGF-1 treated MSC exhibited similar Akt signal modulation via AZ7550 mesylate and GSK1904529A. These observations suggest that IGM-3 potentially activates Akt signaling through similar mechanisms as soluble IGF-1. Compared to cRGD alone, the bifunctional cRGD/IGM-3 hydrogel elicited a marked increase in ERK1/2 phosphorylation, suggesting an additive or synergistic signaling response upon concurrent engagement of both receptors (Fig. 2H).

### Modulation of Cytokine Production in MSC Encapsulated in cRGD/IGM-3 Alginate during Inflammatory Challenge

We first assessed the potential of our three candidate peptides to modulate cytokine provoked MSC secretory behavior (Fig. 3A). IL-1β is present in inflammatory environments into which MSCs are typically transplanted for regenerative applications [33], and can provoke a pro-inflammatory phenotype in MSC [34]. Consistent with this prior observation, IL-1β induced marked secretion of inflammatory cytokines by MSC during culture (Fig. 3B-F). IL-1β increased the secretion of cytokines regardless of hydrogel condition (Fig. 3B-F). However, MSC encapsulated into gels coupled with cRGD and IGM-3 secreted substantially lower levels of TNF-α and granulocyte-macrophage colony-stimulating factor (GM-CSF) compared to MSC cultured in alginate without peptide modifications (“naked” alginate), or alginate modified with cRGD alone (Fig. 3B-C). Furthermore, MSCs cultured within cRGD/IGM-3-alginate secreted the highest levels of Tissue Inhibitor of Metalloproteinases (TIMP 1, 2 and 4) (Fig. 3D-F). This suggests the potential that IGM-3 not only blunts IL-1b-induced inflammation in MSC, but that the MSC under these conditions might secrete factors to attenuate ECM degradation in the context of injuries.

**Figure 3:**
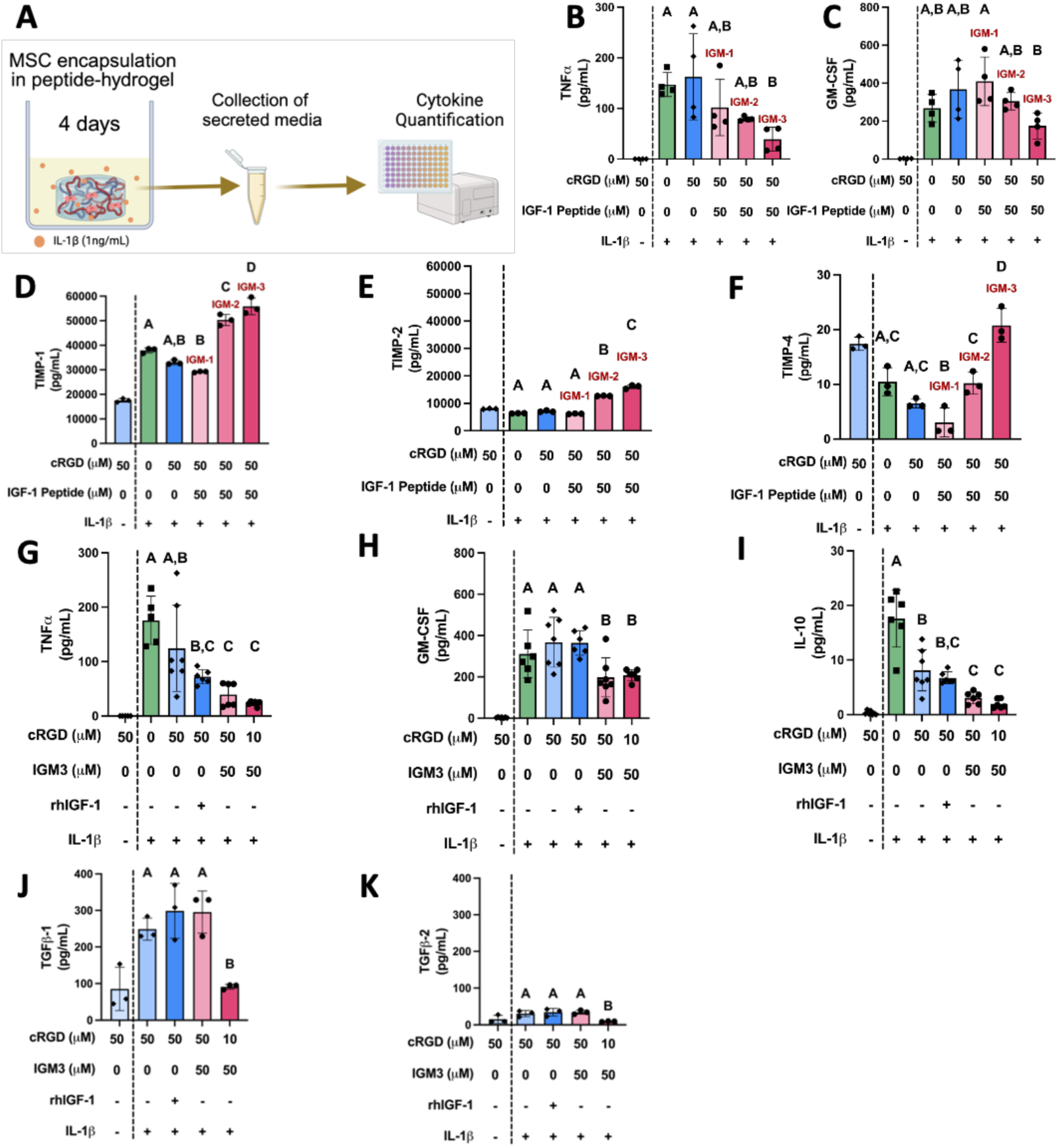
Bioactivity of Insulin-like Growth Factor Peptide Mimetic Challenged with Interleukin-1 Priming. A) Schematic of the experimental setup used to collect secreted medium from MSC hydrogel cultures for quantification. B-C) Quantification of total protein secretion in MSC supernatants. MSCs were encapsulated in various hydrogel conditions and exposed to 1 ng/mL IL-1β for 4 days. Each panel shows the response for different IGM peptide-containing hydrogels (IGM-1, IGM-2, IGM-3). IL-1β priming increased cytokine secretion in all hydrogel conditions, with cRGD and IGM-3 showing a notable reduction in inflammatory cytokines (TNF-α, GM-CSF). D-F) IL-1β increased tissue inhibitor of metalloproteinase (TIMP-1, TIMP-2, and TIMP-4) secretion in MSCs across all hydrogel conditions, with cRGD/IGM-3 showing significant production. G-H) Cytokines associated with inflammation were assessed in MSCs encapsulated in alginate, cRGD-alginate, cRGD-alginate supplemented with soluble IGF-1 (500 ng/mL, rhIGF-1), or various densities of cRGD/IGM-3 after 1ng/mL IL-1β challenge. Secreted pro-inflammatory cytokine levels (TNF-α, GM-CSF) significantly increased after IL-1β priming. The dual-peptide hydrogel containing cRGD and IGM-3 significantly lowered secreted TNF-α and GM-CSF levels compared to controls. I) IL-1β priming led to an increase in the secretion of IL-10. IL-10 secretion varied significantly between cRGD-only alginate and other peptide densities of cRGD/IGM-3. J-K) Secreted TGFβ-1 and TGFβ-2 levels were lower in the 10 μM cRGD/50 μM IGM-3 hydrogel compared to the 50 μM cRGD-only hydrogel. Data represent biological replicates; error bars denote ±SD. Statistical analysis was performed using one-way ANOVA with Tukey’s multiple comparison test; different letters indicate significant differences at *p < 0.05*.

To further explore the immunomodulatory effects of the IGM-3 peptide mimetic under inflammatory conditions, we quantified a broad array of secreted factors linked to pro-inflammatory and anti-inflammatory responses for MSC cultured in the gels presenting cRGD/IGM-3 at the combination yielding optimal survival (Fig. 2B, 4A). As expected, IL-1β challenge increased MSCs’ secretion of pro-inflammatory cytokines, regardless of the alginate hydrogel used for cell encapsulation (Fig. 3G-K, Supp. Fig.2 A-E). Notably, however, MSCs’ secretion of TNF-α and GM-CSF were significantly reduced when these cells were encapsulated into cRGD/IGM-3-alginate compared to unmodified alginate or cRGD-alginate (Fig. 3G-H). This reduction in TNF-α and GM-CSF production was similar to the reduction achieved through exposure of MSC to high doses of soluble IGF-1 in the presence of IL-1β (Fig. 3G-H). IL-10 secretion by MSC dramatically increased in unmodified alginate but decreased in both peptide density of cRGD/IGM-3-alginate ; the upregulation of IL-10 with IL-1β may be consistent with the cytokine’s reported role as regulating both pro and anti-inflammatory responses (Fig. 3I; [35]). In contrast to these findings, neither IGM-3 modified gels nor soluble IGF-1 significantly reduced MSCs’ secretion of IL-6, IL-8, IFNγ, IL-17A, or IL-13 (Supp. Fig.2 A-E).

In addition to inflammation, MSC have also been used to suppress tissue fibrosis, in part by reducing tissue-endogenous TGFβ-1 and TGFβ-2 [36]. Consistent with an ability of inflammatory cytokines to induce a pro-fibrotic phenotype in stromal cells like MSC, both TGFβ-1 and TGFβ-2 secretion were markedly upregulated in alginate-encapsulated MSC when treated with IL-1β. However, the encapsulation in the optimal coupling of cRGD/IGM-3-alginate modestly blunted production of these TGF-β isoforms (Fig. 3J-K); this effect may be more strongly linked to the presence of cell adhesive cRGD as reducing cRGD levels in the alginate gel from 50mM to 10µM reduced TGFβ secretion even with IGM-3 levels held constant (Fig. 3J-K).

### IGM-3 Improves MSCs’ Ability to Diminish Inflammation in Co-cultured Human Intervertebral Disc (IVD) Cells

MSCs exert paracrine effects by secreting growth factors and cytokines that enhance recipient tissue regeneration and cellular survival, while dampening immune responses [37]. Thus, we designed a Transwell-based co-culture study where the secretome of gel-encapsulated MSC was exposed to primary cells derived from human degenerative IVD, where both cell populations were challenged with IL-1β. Degenerative IVDs produce a host of pro-inflammatory cytokines, most notably TNF-a and IL-1β [38, 39] and MSC based therapies have long been evaluated in the treatment of this cascade [40]. To obtain a global understanding of how the MSC secretome might ameliorate this pro-degenerative cascade, we profiled these primary human IVD cells using RNA sequencing (Fig. 4A). Analysis showed that while the MSC-secretome can shift global transcription in target IVD cells when MSC were encapsulated into any alginate hydrogel formulation. However, the combined presence of cRGD and IGM-3 in MSC-encapsulated gels had the most dramatic effect on IVD cells (Fig. 4B-E). This was visualized with Principal Components Analysis (PCA), which showed increased separation between IVD cells exposed to IL-1β without co-culture, versus those exposed to IL-1β in the presence of cRGD/IGM-3-alginate-encapsulated MSC (Fig. 4B). Notably, primary IVD cell populations exposed to IL-1b clustered similarly regardless of whether they were co-cultured with MSC in unmodified alginate gel or MSC in cRGD-alginate. These effects were robust across primary disc cells isolated from 6 patients (3 male, 3 female) who spanned a broad age range (40-70 y.o.). Overlaps in PCA space suggest similar modulatory effects of MSCs in alginate and cRGD alginate on the target disc cells. However, primary disc cells co-cultured with MSC within gels modified with the optimized cRGD/IGM-3 combination clustered distinctly in PCA space. (Fig. 4B).

**Figure 4:**
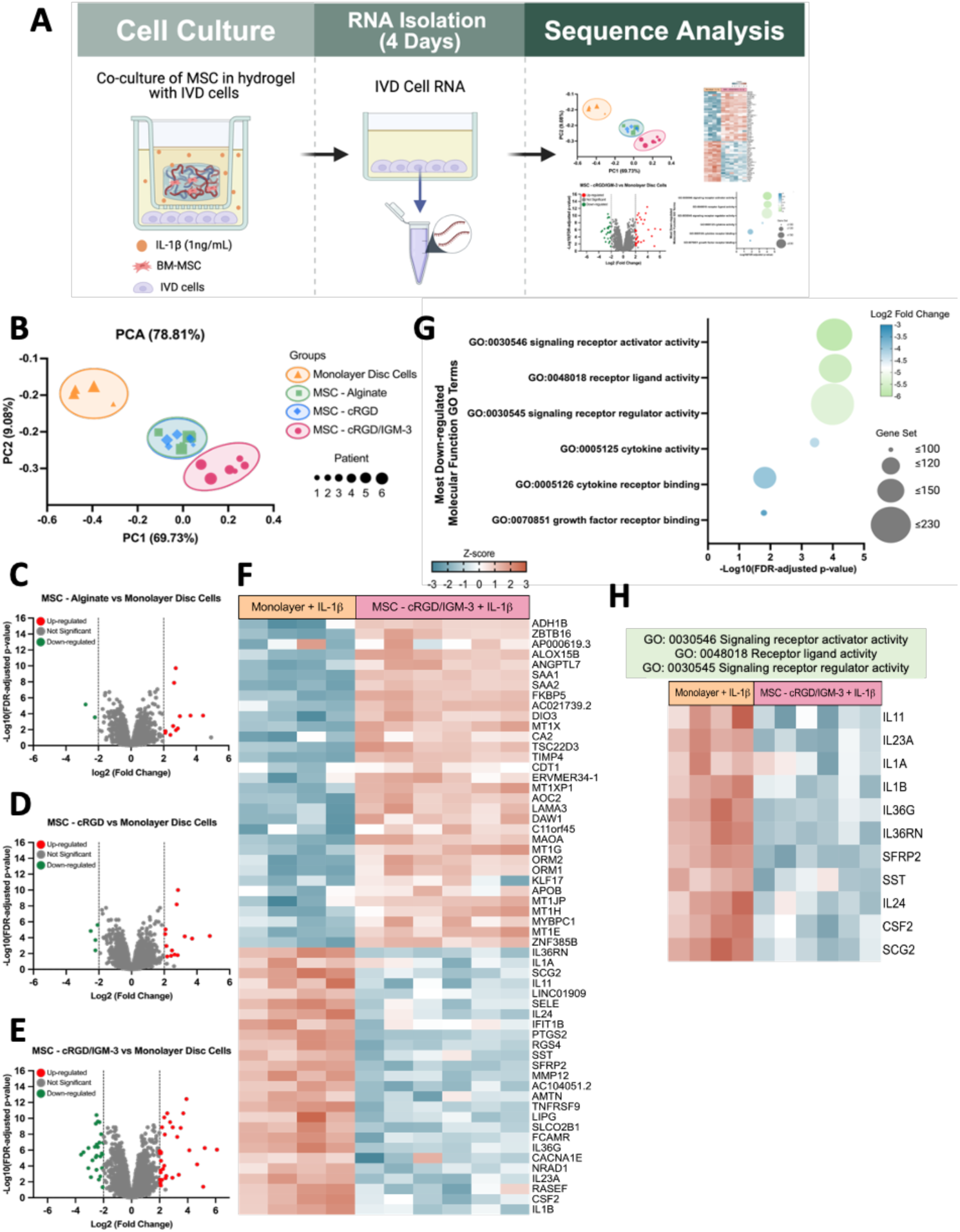
Differential Regulation of Inflammatory Genes in Degenerative Primary Intervertebral Disc Cells Co-Cultured with MSCs in cRGD/IGM3. A) Flow chart depicting bulk RNA-sequencing study. B) Principal Component Analysis (PCA) clearly separates sample groups based on variances: monolayer disc cells (n=4), disc cells co-cultured with MSCs in unmodified alginate (MSC-Alg, n=6), in cRGD alginate (MSC-cRGD, n=6), and in cRGD/IGM3 alginate (MSC-cRGD/IGM-3, n=6). Overlaps in PCA space indicate similar modulatory effects of MSCs in alginate and cRGD alginate on disc cells. C-E) Volcano plots highlight genes differentially regulated in disc cells co-cultured with MSCs in various alginates (MSC-alg, MSC-cRGD, MSC-cRGD/IGM3) compared to monolayer culture, using a cutoff of FDR-adjusted *p-value* < 0.05 and a log2 fold change > 2 or < -2. F) Heatmap of z-score normalized gene expression shows patterns of down-regulated and upregulated genes in disc cells co-cultured with MSC-cRGD/IGM3 versus monolayer. G) Gene Ontology (GO) analysis of the most down-regulated molecular function terms, with a cutoff of log2 fold-change > 2 or < -2 and adjusted FDR *p-value* < 0.05. H) Heatmaps display gene expression profiles mapping to the top three most down-regulated GO terms in the cRGD/IGM-3 condition.

Volcano plot analysis revealed that transcriptional shifts identified in PCA corresponded with a more pronounced differential gene expression in IVD cells co-cultured with MSCs encapsulated in alginate grafted with the optimized combination of cRGD/IGM-3 compared to those in unmodified alginate or cRGD-alginate. This suggests that secreted factors from MSCs in the dual-peptide condition induced significant global phenotypic changes in the IVD cells, particularly in an inflammatory environment (Fig. 4C-E). A heatmap using z-score normalized expression displayed a consistent and distinct pattern of downregulated genes related to inflammation (*adjusted p-value < 0.05*, fold change > 2 or < -2), highlighting the significant immunomodulatory impact of factors secreted by MSCs within the cRGD/IGM-3 hydrogel (Fig. 4F). This modulation is likely important for enhancing tissue repair mechanisms and controlling inflammatory responses suggesting that MSCs influence not only individual cell behavior but also the broader tissue environment [41]. This finding highlights MSCs’ paracrine effects in actively modulating the inflammatory response of cells from a disease state at a transcriptional level. By downregulating pro-inflammatory genes, MSCs exerted a paracrine influence that steers the immune environment towards a reparative state, with potential to enhance tissue repair and attenuate progressive degeneration.

Gene Ontology (GO) analysis provided insights into the molecular function processes that were differentially regulated in IVD cells co-cultured with MSCs in cRGD/IGM-3. Notably, the terms ‘signaling receptor activator activity,’ ‘receptor ligand activity,’ and ‘signaling receptor regulator activity’ were among the top three most significantly regulated by the experimental condition (*adjusted p-value < 0.05*, fold-change > 2 or < -2; Fig. 4G). A heatmap using z-score normalized expression for the top three GO terms displayed a consistent pattern of downregulated inflammation-related genes (*adjusted p-value < 0.05*, fold change > 2 or < -2), such as IL11, IL23A, IL-1β, and IL1A, and CSF2 (Fig. 4H). This downregulation further highlights the immunomodulatory effect of factors secreted by MSCs within the cRGD/IGM-3 hydrogel. The prominence of these changes suggests that the MSCs, through the cRGD/IGM-3 hydrogel, play a pivotal role in modulating cell signaling pathways critical for cellular communication and response to environmental cues.

## Discussion

Recent works have shown phage-derived peptides can bind specifically to stem cell receptors, enabling their use in functionalizing synthetic scaffolds for stem cell expansion *in vitro* [42]. These peptides have been incorporated into biomaterials to bind to adhesive or growth factor receptors and shown to promote cell growth and functionality [43, 44]. Our findings demonstrate that the IGM-3 peptide sequence, discovered through phage-display, supports MSC survival and exerts significant immunomodulatory effects in a serum-free environment while two other sequences, IGM-1 and IGM-2 exhibit some but lesser bioactive effects upon MSCs. Importantly, although IGM-3 lacks sequence similarity to natural IGF-1, this peptide mimetic could activate downstream signaling similarly to soluble IGF-1 (Fig. 2C), supporting cell survival and cytokine production of MSCs in the presence of IL-1β. Studies with IGF-1 receptor and insulin-receptor specific small molecule inhibitors and IGF-1 receptor antibody blocking suggest that the mechanism of action for IGM-3 is similar to that of IGF-1 (Fig. 2D-G). This suggests that despite lacking homology to full-length IGF-1, IGM-3 engages the IGF-1 receptor to activate downstream pathways in a manner analogous to the native IGF-1 ligand to modulate MSC function. The C-domain of IGF-1 (IGM-1) has been previously studied for its role in cell survival and anti-inflammatory signaling in MSCs [21, 22]; however, IGM-3 proved to be more potent for both applications in the present study (Fig. 2A; Fig. 3B & C; Fig. 3G & H). IGM-2 also showed less potency compared to IGM-3 (Fig. 2A), which could be due to its selective binding to the insulin receptor as shown by other studies [45] since insulin receptor is to mediate the metabolic effects of insulin, whereas the IGF-1 receptor is involved in mitogenic signaling. The selective binding of each of the IGM peptide mimetic suggests a different profile with regard to cellular signaling and mitogenic activity.

IGF-1 receptor signaling is well-documented for its critical role in promoting cell survival, proliferation, and immunomodulation through the PI3K/Akt pathway [28, 46, 47]. Our observation also points to a role for IGF-1-induced signaling in immunomodulation in which the addition of soluble IGF-1 or IGM-3 peptide mimetic in combination of cRGD lowered the production of inflammatory cytokines (Fig. 3G). Previous research indicates that IGF-1 can suppress the expression of toll-like receptor 4 in skeletal muscle cells, reducing inflammation via the PI3K/Akt pathway and consequently lowering TNF-α and IL-6 levels [48]. In addition to its effects through the PI3K/Akt pathway, IGF-1 has been shown to modulate MSCs by activating the JAK/STAT pathway, thereby enhancing their survival and anti-inflammatory capabilities [49, 50]. For example, Takahashi *et al.* [51] and colleagues elucidated the activation of JAK/STAT signaling pathways by IGF-1 in rat cardiomyocytes, in which there is a complex interplay of kinase activities and phosphorylation events essential for cellular responses to growth signals. Based on these prior results, it is plausible that culturing MSC within the cRGD/IGM-3-hydrogels, which markedly reduced both pro-inflammatory cytokines (e.g. TNF-a, Fig. 3G), but also variably affected anti-inflammatory cytokines (e.g. IL-10; Fig. 3I), activated JAK/STAT to antagonize inflammatory signaling pathways. Interestingly, co-engagement of integrin α_v_β_3_ by cRGD and IGF-1R by IGM3 may produce a combinatorial or synergistic enhancement of both MAPK and PI3K/Akt signaling. For example, in vascular smooth muscle cells, pharmacological blockade of α_v_β_3_ markedly attenuates IGF-1 driven ERK1/2 and Akt phosphorylation, thus inhibiting proliferation and migration [52, 53]. More broadly, integrin mediated adhesion to extracellular matrix proteins has been shown to amplify growth factor receptor outputs [54], and attachment through α_v_β_3_ potentiates Ras/MAPK and PI3K/Akt activation beyond levels seen with receptor tyrosine kinase ligation alone in endothelial and stromal cells [55]. Our observation that dual MSCs cultured on cRGD/IGM3 hydrogel markedly elevates ERK1/2 phosphorylation (Fig. 2H) while maintaining robust Akt signaling (Fig. 2C); therefore, suggests a synergistic cross-talk between integrin and IGF-1R inputs. Altogether, these findings suggest that IGM-3 not only mimics IGF-1 signaling but may amplify regenerative signaling through integrin-mediated synergy.

The therapeutic efficacy of MSCs in exerting trophic effects on endogenous cells upon transplantation within inflammatory environments is predominantly attributed to paracrine signaling rather than direct cell integration [37, 56, 57]. Our findings revealed that regardless of the micro-environment that MSC were encapsulated into (e.g. unmodified alginate vs. cRGD vs. cRGD/IGM-3), the secretome of these cells strongly affected global RNA expression of primary cells derived from degenerative human IVD that were provoked with IL-1β (Fig. 4C-E). However, the most dramatic RNA expression shifts were observed in disc cells co-cultured with MSC encapsulated into cRGD/IGM-3-alginate (Fig. 4C-E). Notably, we observed a downregulation of genes such as TNFRSF9, CSF2, and IL-1β, which are associated with anti-inflammatory responses, suggesting a shift towards a less inflammatory state in IVD cells co-cultured with MSCs within the cRGD/IGM-3-alginate (Fig. 4F). These paracrine effects of MSCs in modulating a targeted endogenous cell population *in vitro* are consistent with prior reports. For example, Levorson *et al.* demonstrated that chondrocyte proliferation and expression of ECM proteins like aggrecan that are associated with healthy cartilage were enhanced when the chondrocytes were co-cultured with MSC [10]. Other studies have demonstrated that MSCs co-cultured with degenerated nucleus pulposus cells of the IVD upregulated TGFβ-1, IGF-1, and aggrecan gene expression in these cells, while MSCs had no effect on the phenotype of non-degenerate nucleus pulposus cells. This suggests that secreted factors by MSCs could stimulate the endogenous IVD cell population to regain a nondegenerative phenotype [58].

Taken together, our study illustrates the significant potential of IGF-1 peptide mimetic and cell adhesive ligand in enhancing the survival and immunomodulatory properties of MSCs. MSC within the cRGD/IGM-3-alginate secreted factors that downregulated inflammatory gene expression in primary human degenerative IVD cells, which are typically highly inflammatory during degeneration. These findings illustrate the design of a bioactive biomaterial to retain and enhance stem cell functionality using IGF-1 peptide mimetics and circumvent a potential need to co-administer the cells together with high doses of soluble growth factors. Future work could focus on exploring the synergism between JAK/STAT, PI3K/AKT, and cell adhesive signaling pathways and cellular mechanisms that influence MSC immunomodulation will offer deeper mechanistic insights into the potential for co-delivery of MSCs and growth factor peptide mimetics in the development of MSC therapies.

## Supporting information

Suppemental Figure NEW

## Acknowledgments

Supported with funds from the National Science Foundation Graduate Research Fellowship Program under Grant No. DGE-1745038 and DGE-2139839, from National Institute of Biomedical Imaging and Bioengineering Training Grant T32 (EB028092), from the National Institute of Arthritis and Musculoskeletal and Skin Diseases (R01AR077678 and R01AR077678-02S1), and from the National Heart and Lung Institute (R01HL159094). We thank the Genome Technology Access Center at the McDonnell Genome Institute at Washington University School of Medicine for help with genomic analysis. The Center is partially supported by NCI Cancer Center Support Grant #P30 CA91842 to the Siteman Cancer Center from the National Center for Research Resources (NCRR). This publication is solely the responsibility of the authors and does not necessarily represent the official view of NSF, NIH or NCRR.

