## Supplementary material for "IGF-1 Peptide Mimetic-functionalized Hydrogels Enhance MSC Survival and Immunomodulatory Activity": Suppemental Figure NEW

### Supporting Information Text

Cell adhesion receptors often transduce signals with other receptors such as IGF-1 or TGF-β receptor to dictate cell fate [1]. As shown in our studies, the addition of cRGD in combination of IGM-3 peptide mimetic is necessary to support MSC survival. Cell viability of MSCs encapsulated in IGM-3 only alginate exhibited significantly lower viability compared to the material presented with both cRGD and IGM-3 peptide mimetic (Fig. S1).

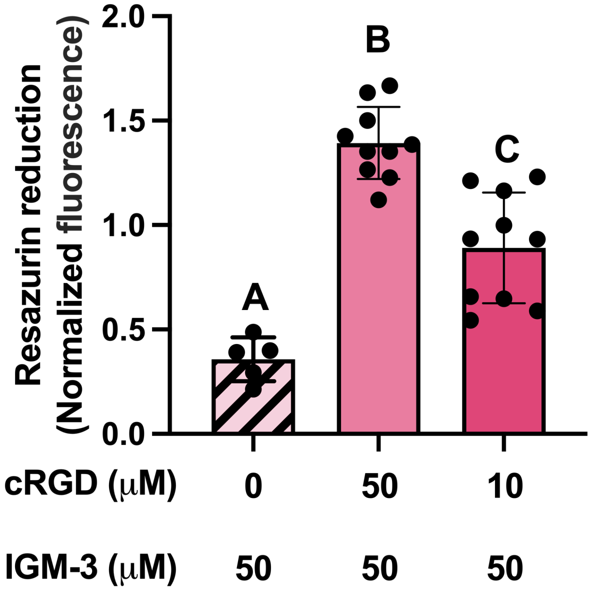


**Supplementary Fig. 1. Combination of cRGD and IGM-3 Peptide Mimetic Supports Cell Survival.** Comparison of resazurin reduction in MSCs encapsulated in IGM-3-alginate or varied peptide density of cRGD and IGM-3 on Day 7, normalized to Day 1 values. Data points represent biological replicates with error bars indicating ±SD. Statistical analyses were performed using one-way ANOVA with Tukey's multiple comparisons test; non-significant results are indicated by the same letter, while significant differences are marked with different letters at $p < 0.05$.

Neither IGM-3 modified gels nor soluble IGF-1 significantly reduced MSCs' secretion of IL-6, IL-8, IFNγ, IL-17A, or IL-13 (Supp. Fig.2 A-E).

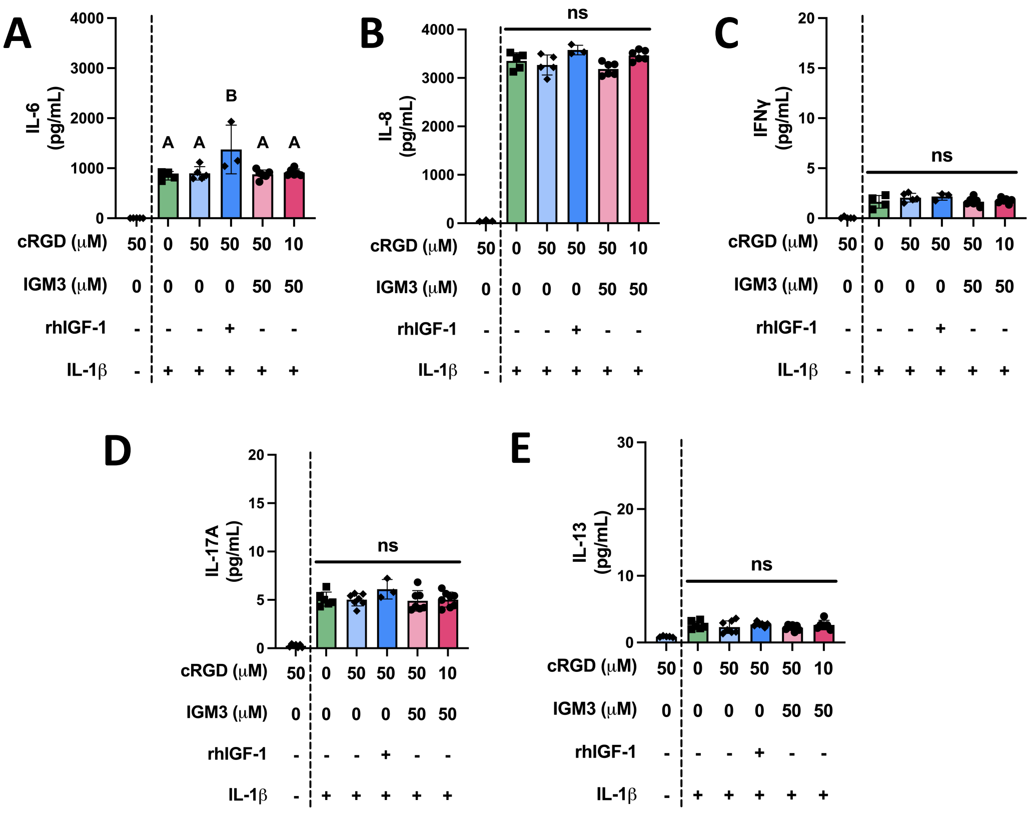


**Supplementary Fig. 2. Interleukin-1β Priming Enhanced both Anti-inflammatory and Pro-inflammatory Cytokine Production in MSCs across all Hydrogel Conditions.** A-D) Cytokines associated with inflammation were assessed in MSCs encapsulated in alginate, cRGD-alginate, cRGD-alginate supplemented with soluble IGF-1 (500 ng/mL, rhIGF-1), or various densities of cRGD/IGM-3 after 1ng/mL IL-1β challenge. Secreted pro-inflammatory cytokine levels (IL-6, IL-8, IFNγ, IL-17A) significantly increased after IL-1β priming. There was no observed difference in IL-8, IFNγ, and IL-17A. IL-6 levels remained consistent across all hydrogel conditions, except for a slight increase in cRGD supplemented with 500 ng/mL rhIGF-1. E) IL-1β priming led to an increase in the secretion of cytokines associated with anti-inflammation. IL-13 production was consistent across all conditions.
